# An Analysis of gRNA Sequence Dependent Cleavage Highlights the Importance of Genomic Context on CRISPR-Cas Activity

**DOI:** 10.1101/2021.05.06.442929

**Authors:** E.A Moreb, M.D. Lynch

## Abstract

CRISPR-Cas9 is a powerful DNA editing tool. A gRNA directs Cas9 to cleave any DNA sequence with a PAM. However, some gRNA sequences mediate cleavage at higher efficiencies than others. To understand this, numerous studies have screened large gRNA libraries and developed algorithms to predict gRNA sequence dependent activity. These algorithms do not predict other datasets as well as their training dataset and do not predict well between species. To better understand these discrepancies, we retrospectively examine sequence features that impact gRNA activity in 39 published data sets. We find strong evidence that the genomic context, which can be defined as the DNA content outside of the gRNA/target sequence itself, greatly contributes to differences in gRNA dependent activity. Context underlies variation in activity often attributed to differences in gRNA sequence. This understanding will help guide future work to understand Cas9 activity as well as efforts to identify optimal gRNAs and improve Cas9 variants.

**Highlights:**

- Species-specific genomic context drives variability in gRNA activity in a PAM proximal sequence-dependent manner
- Increased PAM specificity of Cas9 and/or increased Cas9/gRNA expression reduces the impact of species-specific context
- Current gRNA prediction algorithms trained on species are not expected to predict activity in another species

## Introduction

Since their discovery in 2012, CRISPR systems have revolutionized how we manipulate biology.^1^ The successful application of CRISPR systems is dependent on the guide RNA (gRNA) but understanding which gRNA sequences effectively cleave their targets has proved challenging.^2–4^ Predictive algorithms have been developed to select gRNA with improved on-target activities.^3,5–10^ These algorithms rely on sequence features of the gRNA. (Figure 1a) While many of these algorithms have achieved good predictability within their training data, predictions between datasets, particularly between different species, are not as accurate.^8,10–15^ This suggests that the features used to develop these algorithms are not effectively capturing changes in context (Figure 1 b-c). Broadly defined, “context” includes all variables outside of the gRNA/Cas9 complex that can impact activity, with the “genomic context” more specifically pertaining to the DNA sequences outside of the gRNA.^16^ While significant work has been done to characterize the factors impacting Cas9 activity *in vitro*, there are limitations when comparing this to *in vivo* activity, particularly with respect to genomic context. The amount of competitive Cas9 binding sites^16^, target site accessibility, and other contextual *in vivo* factors are not easily replicated *in vitro.*

**Figure 1:**
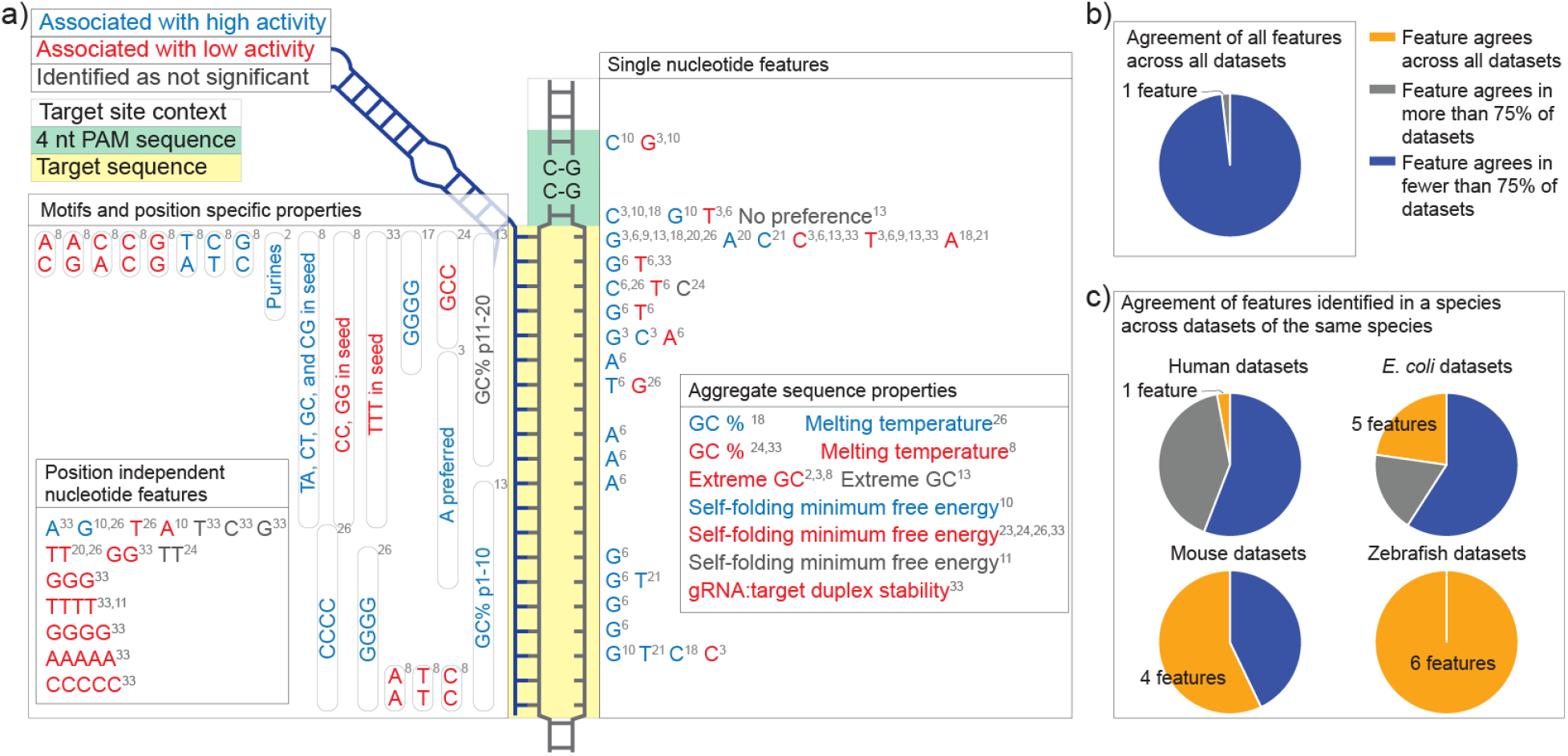
Summary of gRNA specific features identified as most important. a) The gRNA:target duplex is highlighted in yellow and the PAM site is highlighted in green. Features that positively impact gRNA activity, have been identified as inhibitory to activity, or were specifically identified as not significant are labeled in blue, red, and grey, respectively. Position dependent features are labeled in the relevant position while position independent features are shown in separately labeled boxes. We then compared feature activity across datasets to determine if the impact of the features was positive or negative across datasets. For discrete features, we calculated a log odds ratio comparing the top third and bottom third of gRNA activity in each dataset. For continuous features, we correlated the features with gRNA activity. The sign of the log odds ratio or correlation was used to determine if a feature agreed across multiple datasets. b) When comparing all datasets, there was no feature identified with a common impact on activity across all datasets and only one feature identified across 75% of the datasets. c) We next looked at what species each feature was identified in and compared the relative impact of that feature across datasets collected within that species. Of features identified as important in human datasets, only 1 feature consistently had the same impact across human datasets. For other species, we identified five, four, and 6 features that were consistent across datasets within the *E. coli*, mouse, and zebrafish datasets. *Y. lipolytica* was excluded from this analysis as there is only one dataset.

The context can greatly affect gRNA activity. For example, 4 thymines in a row in a given gRNA is a termination sequence (leading to low gRNA expression) in some contexts (Supplemental Figure S1).^17^ Another example is the inhibitory search space (transient binding to “non-target sites”), part of the genomic context, which we have recently reported.^16^ We demonstrated that the efficiency with which Cas9 cleaves a target site is decreased by the addition of inhibitory “non-target” sequences which are transiently interrogated by the Cas9/gRNA complex but not cleaved.^16^ In the present study we sought to better understand the impact of context on gRNA activity and toward this goal we report a retrospective analysis of 39 gRNA library datasets from different species, with several Cas9 variants, using both endogenous and exogenous target sites, and in different experimental systems.^3–11,16,18–26^ In this analysis we confirm the importance of context as a key factor driving gRNA dependent activity *in vivo*.

## Results

We began by compiling data as discussed in the Methods Section, and illustrated in Figure 2a.^3–11,16,18–26^ The datasets have varied distributions of cutting/cleavage activity, from binary distributions (gRNAs that either cut or do not cut) to skewed or normal distributions, suggesting significant experimental and context dependent differences in gRNA dependent activity (Figure 2b, data compiled in Supplementary File 1). Despite these differences, most datasets accurately capture the reported four nucleotide PAM preference of Cas9, highlighting that Cas9 specific features should correlate across contexts (Supplemental Figure S2). We therefore sought to better understand how gRNA sequence specific activity is impacted by context.

**Figure 2:**
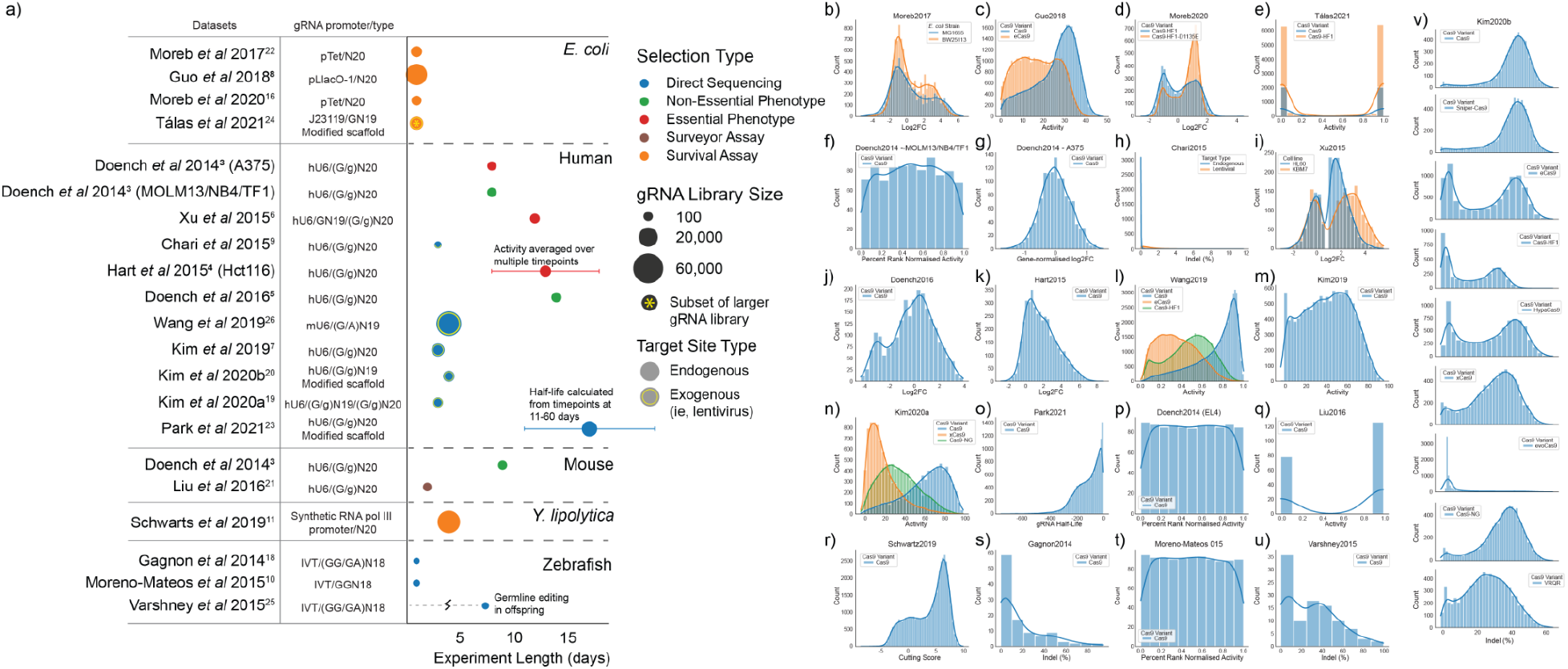
Summary of collected datasets. a) Datasets collectively represent a diverse set of expression methods, experiment durations, species, selection types, gRNA library sizes, and target types. b-v) These datasets cover a broad range of activity distributions, including moderate to extremely binary distributions, normal distributions, and completely uniform distributions, based on how activity was measured and data were processed. All datasets are shown with higher gRNA activity represented by larger numbers. In some cases, that required inverting the scale of activity (namely, b, d, and o).

### PAM proximal sequence is most predictive of Cas9 activity

All on-target prediction algorithms heavily rely on gRNA sequence to predict activity. We therefore sought to understand how predictive the full gRNA sequence is, as well as which part of the sequence is most predictive within each dataset. Previously, gRNA sequences have been used as features in algorithms by using one hot encoding of overlapping dinucleotides.^5^ We therefore used the same approach to digitize all sequences (Figure 3a). For each dataset, after converting gRNA sequences to a one hot matrix encoding dinucleotides, we randomly split the dataset into a training group and testing group representing 80% and 20% of the gRNA, respectively. After training, we predicted the activity in the test group and compared the predicted activity to actual activity using a Pearson correlation. We performed 10-fold cross validation by splitting the training and test groups randomly each time and averaging the results (Figure 3b). In small datasets with n < 205 gRNA, this approach did not prove to be predictive. Similarly, in the data from Chari *et al* 2015, the activity for gRNA targeting endogenous sites was not predictable, likely due to the low overall activity within this dataset (Figure 2h). Among the remaining datasets, the Pearson values ranged from 0.18 to 0.84 highlighting both the link between sequence and activity and the variability of sequence impact in different experimental contexts.

**Figure 3:**
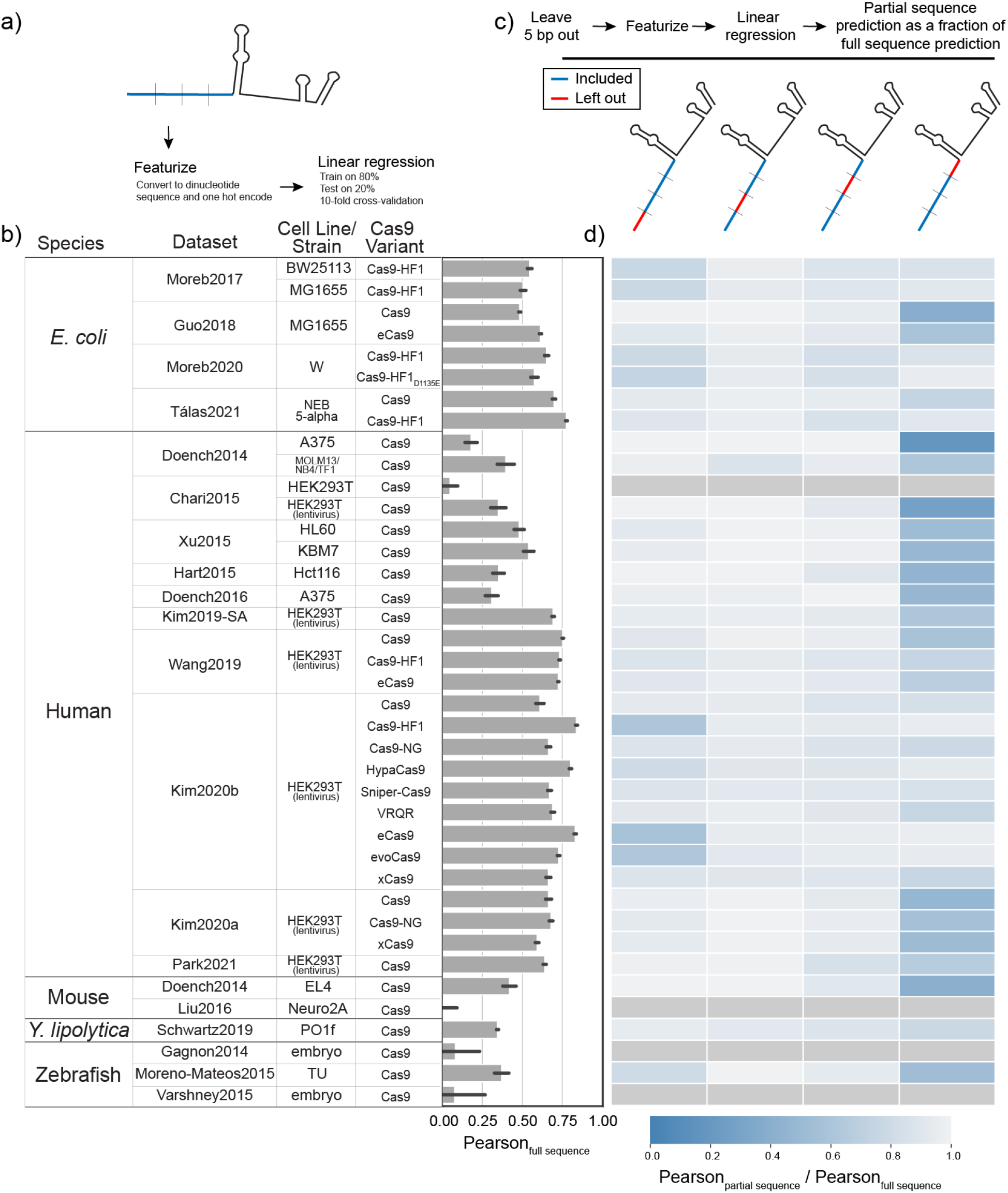
The PAM proximal portion of the gRNA provides most of the predictive power of the sequence. a) To better understand gRNA sequence based predictions, the 20 bp targeting sequence of each gRNA was converted to a dinucleotide one hot matrix and used to predict activity with a linear regression. For each dataset, 80% of gRNA were randomly assigned to a training group while the remaining 20% were used as a test group. Predicted activity was compared with actual activity using Pearson correlation coefficient and this process was repeated 10 times to achieve 10-fold cross validation. b) The average of the 10-fold cross validation is shown for each dataset. c) We then repeated this analysis but left out 5 bp at a time. d) The heatmap shows the averaged Pearson with 5 bp left out (Pearson_partial sequence_) as a fraction of the averaged Pearson using all 20bp (Pearson_full sequence_). In cases where the Pearson_full sequence_ average was close to zero, we excluded these datasets from further analysis.

We next proceeded to iteratively repeat the linear regressions, each time removing one quarter of the gRNA sequence and correlating the remaining sequence with activity (Figure 3c). The Pearsons for the partial sequence predictions as a fraction of the Pearson for full sequence prediction are given in Figure 3d. These results highlight the majority of the predictive ability of the full gRNA sequence is from the PAM proximal region. The PAM distal 5 base pairs is also important for some datasets, primarily those utilizing high-fidelity variants of Cas9. This result is consistent with the PAM proximal features identified in the literature (Figure 1a).

### The Impact of the PAM proximal sequence on Cas9 Activity is context dependent

To better understand these results, we next examined specific sequence preferences within each dataset. One way to better understand this sequence preference is to compare preference between species. If nucleotide preference is derived from Cas9 itself, we would expect to see strong agreement on sequence preference between datasets, regardless of species.^2^ Intraspecies correlation with a reduced correlation between species would suggest that differences are driven by the larger genomic sequence or context, potentially due to different inhibitory non-target site pools.^16^ The lack of any intra or interspecies correlation would suggest other confounding and unknown context dependent factors.

To investigate the species specific sequence preference in the PAM proximal position, we first determined what length of sequence to compare. In each dataset, we first measured the fractional representation of all possible k-mers (length 1 to 10) starting at the PAM proximal position (Figure 4a). With the exception of the dataset from Hart *et al* 2015, all datasets contained gRNA representing all 16 possible dinucleotide sequences in the PAM proximal position. In Hart *et al* 2015, the dataset was designed to exclude thymines in the four PAM proximal positions, explaining the lack of specific dinucleotide sequences in this dataset.^4^ We grouped gRNA within the remaining datasets by the PAM proximal dinucleotide sequence, calculated the average activity for each group, and then looked at the correlation between these dinucleotide group averages between datasets, in a pairwise-fashion, as demonstrated in Figure 4b (see Supplemental Figure S3 for grouped averages per dataset). In these results, we see low interspecies correlations, but strong intraspecies correlations within the *E. coli*, human, and zebrafish datasets (Figure 4c). In *E. coli*, there are strong correlations between our two previously reported datasets and that of Guo *et al* 2018 but weak to no correlation with the datasets from Talas *et al* 2021. This study used an experimental design, including extrachromosomal targets, enabling rapid gRNA cleavage. As a result, in Tálas *et al* 2021, gRNA are mostly inhibited by the formation of unwanted secondary structures that render gRNA unable to bind the target site (the authors note predicted minimum free energy of gRNA secondary structure is strongly correlated with their library). Within the two mouse datasets, we don’t see a good correlation but this is consistent with earlier results suggesting that the data reported by Liu *et al* 2016 is not large enough and does not have high enough resolution to capture key sequence features driving activity. In *Y. lipolytica*, with only one dataset, we can only conclude that this dataset is not strongly correlated with other species, which agrees with the authors findings that several previously published predictive algorithms for both human and *E. coli* gRNA had no predictive ability on their dataset.^11^ Similarly, within the three zebrafish datasets there is strong correlation when comparing data from Moreno-Mateos *et al* 2015 with the other two but no correlation between the smaller datasets. Taken together, low interspecies correlations, but strong intraspecies correlations, strongly suggest species dependent differences between gRNA activities and a role for the larger genomic context in determining gRNA sequence dependent activity.^16^

**Figure 4:**
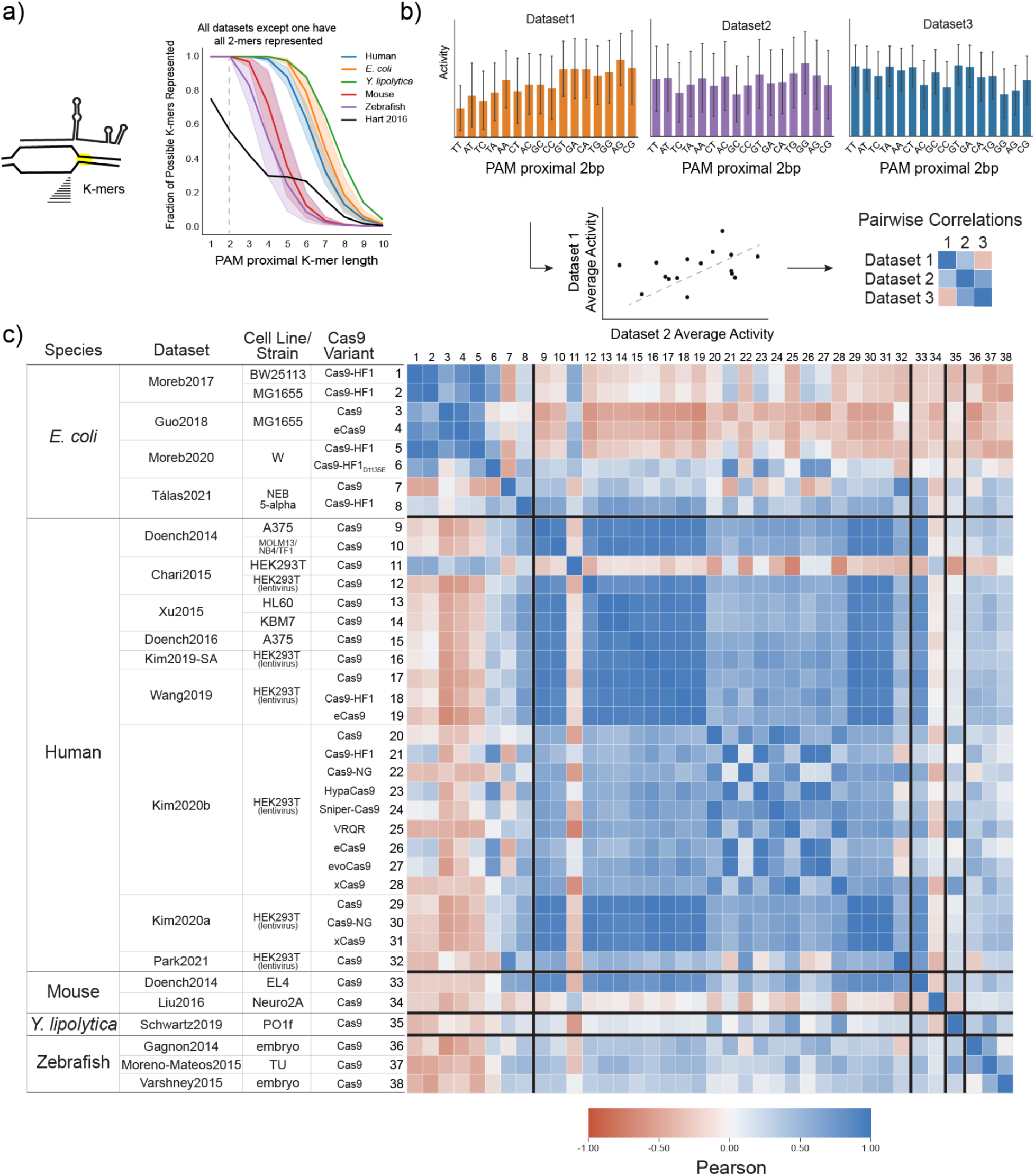
PAM proximal sequence preference is context dependent. a) To understand the sequence preference of the PAM proximal portion of the gRNA, we first determined the best length of PAM proximal k-mers to compare. We looked at the fraction of possible k-mers for each length, k, starting in the PAM proximal position and show that 2-mers are represented in all datasets, except Hart *et al* 2015 which excluded thymines from the PAM proximal 4 bases. We therefore excluded Hart *et al* 2015 from this analysis. b) We next grouped gRNA within each of the remaining datasets by the PAM proximal dinucleotide and calculated the average activity for each dinucleotide group. These averaged values were then correlated in a pairwise fashion between datasets to determine the similarity of dinucleotide sequence impact at this position. c) The heatmap shows Pearson correlations between the averaged values for PAM proximal dinucleotides in all datasets, with blue being more positively correlated and red being more negatively correlated. Datasets are grouped by species and then ordered by year of publication. See Supplemental Figure S3 for comparison of individual datasets.

Within species, several factors appear to reduce intraspecies correlations, suggesting a reduced impact of host context on activity. In *E. coli* datasets, the addition of the D1135E mutation^27^ to Cas9-HF1 appears to change PAM proximal sequence preference, as evidenced by reduced correlation with other *E. coli* datasets (in contrast to differences between high fidelity variants and wild-type Cas9, Supplemental Figure S3). As we previously reported, reducing the search space of Cas9 by increasing PAM specificity, thus reducing potential interactions at non-target sites, resulted in higher overall on-target activity. Another potential approach to reducing the context dependence of gRNA activity is highlighted by the weaker correlations of Kim *et al* 2020b datasets and Park *et al* 2021 dataset with other human datasets. Both of these datasets utilize different sgRNA scaffolds that modify the four consecutive thymines present towards the 5’ end of the scaffold.^20,23^ As four thymines in a row is a known transcription terminator for the eukaryotic RNA polymerase III, the result of this modification is higher expression of the gRNA.^28^ This suggests that increased gRNA expression reduces the impact of context on activity.

### For a given genomic context, the PAM proximal sequence correlates with an upper limit of gRNA activity

We next looked at how well a longer PAM proximal sequence correlates with activity among the human datasets. We previously looked at the fractional representation of all k-mers (length 1 to 10) in the PAM proximal position (Figure 4a). From this analysis, we selected the two largest human datasets (from Wang *et al* 2019 and Kim *et al* 2019) that used the conventional gRNA scaffold in order to have full representation of all possible 5-mer sequences (Figure 5a). We combined these datasets, grouped gRNA by their PAM proximal 5 bp sequence, calculated the average activity for each group and then used this averaged activity to predict gRNA activity in all human datasets based on the PAM proximal 5bp for each gRNA (Figure 5b). Upon correlating predicted activity with actual activity, we found that this approach was reasonably predictive for datasets using the conventional gRNA scaffold while less predictive of gRNA with a modified gRNA scaffold, in line with earlier analysis (Figure 5c). We also confirmed that grouping by the PAM proximal 5 bases was more predictive than grouping by fewer nucleotides (Supplemental Figure S4). Notably, with the exception of high fidelity variants and datasets with modified gRNA scaffolds, these predictions are comparable or better than our earlier linear regression-based predictions using the full gRNA sequence. We then compared predicted activity to actual activity for each dataset (Figure 5d). Again, this comparison highlights the predictive power on datasets using the conventional gRNA scaffold as compared to modified scaffolds. Interestingly, these results illustrate that while this prediction often generates false positives, it generates far fewer false negatives (Figure 5e-g) thus suggesting that given a particular genomic context, the PAM proximal sequence correlates with an upper bound of the potential activity of a given gRNA.

**Figure 5:**
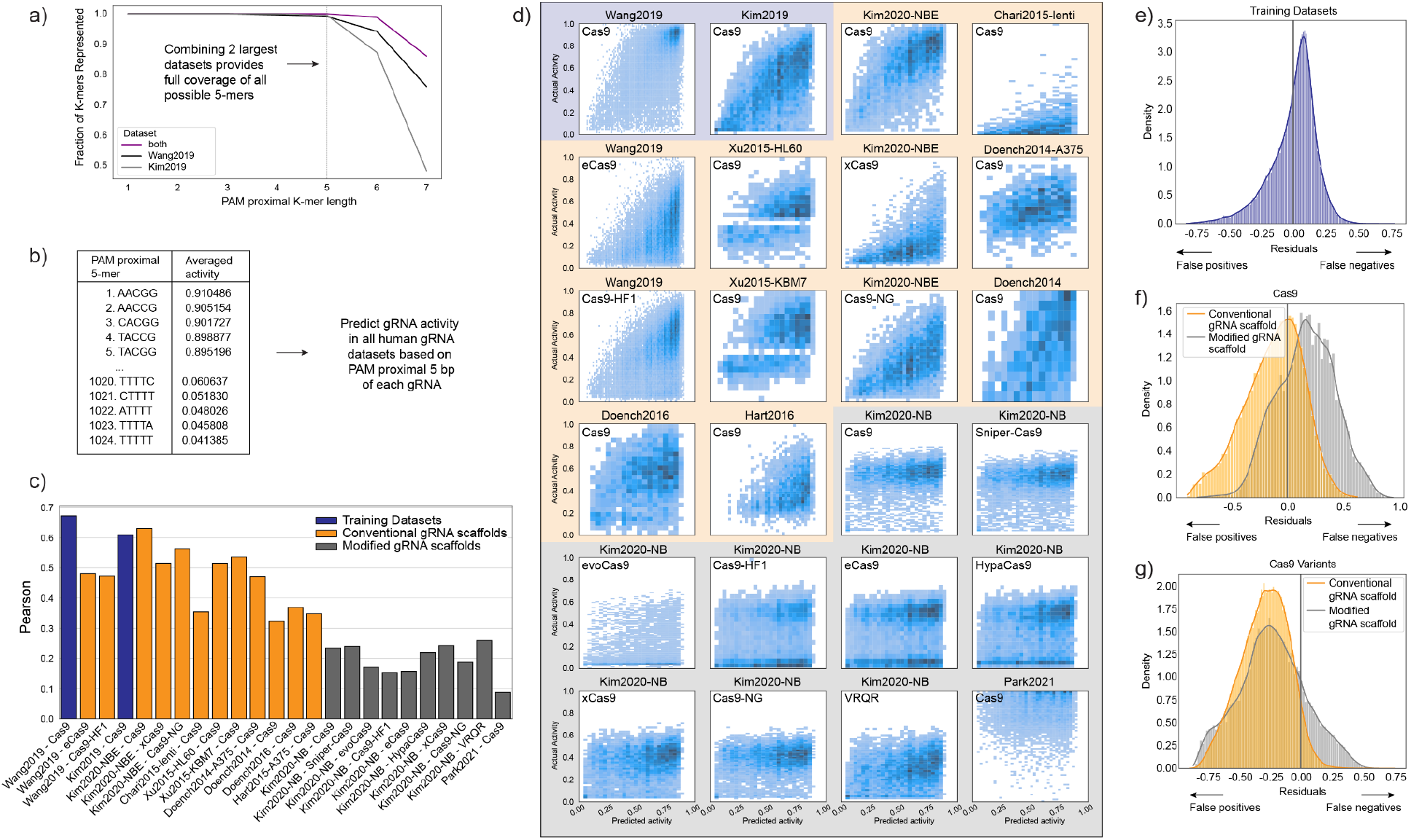
Within human datasets, the PAM proximal sequence is predictive of activity. a) Together, the two largest human datasets contain gRNAs that represent all possible 5-mers in the PAM proximal position. b) We combined these datasets, grouped all gRNA by the PAM proximal 5 bp, calculated an average value for each group, and then used these grouped averages to predict gRNA activity in all human datasets. c) We correlated this predicted activity with actual activity using Pearson correlation. The datasets that we used to generate the averages are highlighted in blue, while test datasets using the conventional sgRNA scaffold or a modified version of the scaffold are highlighted in orange and grey, respectively. d) For each of these datasets, we compared predicted activity on the x-axis to actual activity on the y-axis. We then calculated the residuals (Activity - Predicted Activity) for e) the two training datasets, f) all of the wild-type Cas9 datasets, and g) all other Cas9 variants. Datasets using the normal gRNA scaffold are in orange and those using the modified gRNA scaffold are in grey. Refer to Supplemental Figure S5 for a similar analysis for *E. coli* datasets.

## Discussion

Genomic context determines activity at least in part through an interaction with the PAM proximal sequence of the gRNA, as demonstrated in Figure 5. Predictions based on the PAM proximal sequence improve by including longer sequences (Supplemental Figure S4). However, within the current human datasets, we are limited to using 5 nucleotides of sequence due to a lack of representation of longer sequences. In the future, better coverage of longer sequences may enable an improved understanding of how context dictates activity. This highlights a gap in current gRNA library designs, which has limited the ability to understand context as a key feature driving CRISPR-Cas activity.

Genomic context can explain variability in gRNA activity across species. As illustrated in Figure 6, species specific algorithms to predict gRNA activity may be useful but predicting between species is not appropriate with current gRNA sequence-based features.^8,10,11,13^ Previously these differences in activity have been attributed to different gRNA/Cas9 expression methods, different mechanisms of repair, target site accessibility, or phenotypic screening versus more direct methods of measuring activity.^8,10,14,15^ Our analysis suggests that while expression levels matters, promoter differences are less likely to be driving differences between species. Similarly, differences in repair or target site accessibility may be impactful but would not explain the differences we observe in the PAM proximal sequence preferences between species. Furthermore, while better algorithms, such as deep learning,^7,29^ may improve species-specific predictions, a better understanding of genomic context will be required to predict activity across species. Understanding the impact of novel contexts on gRNA sequence dependent activity is key to developing CRISPR-based applications in new organisms, where current datasets are not expected to be predictive (Figure 6e). To aid these future efforts, we have provided a proposed workflow and key considerations when designing gRNA libraries to better develop gRNA design algorithms in new systems (Supplemental Note 1).

**Figure 6:**
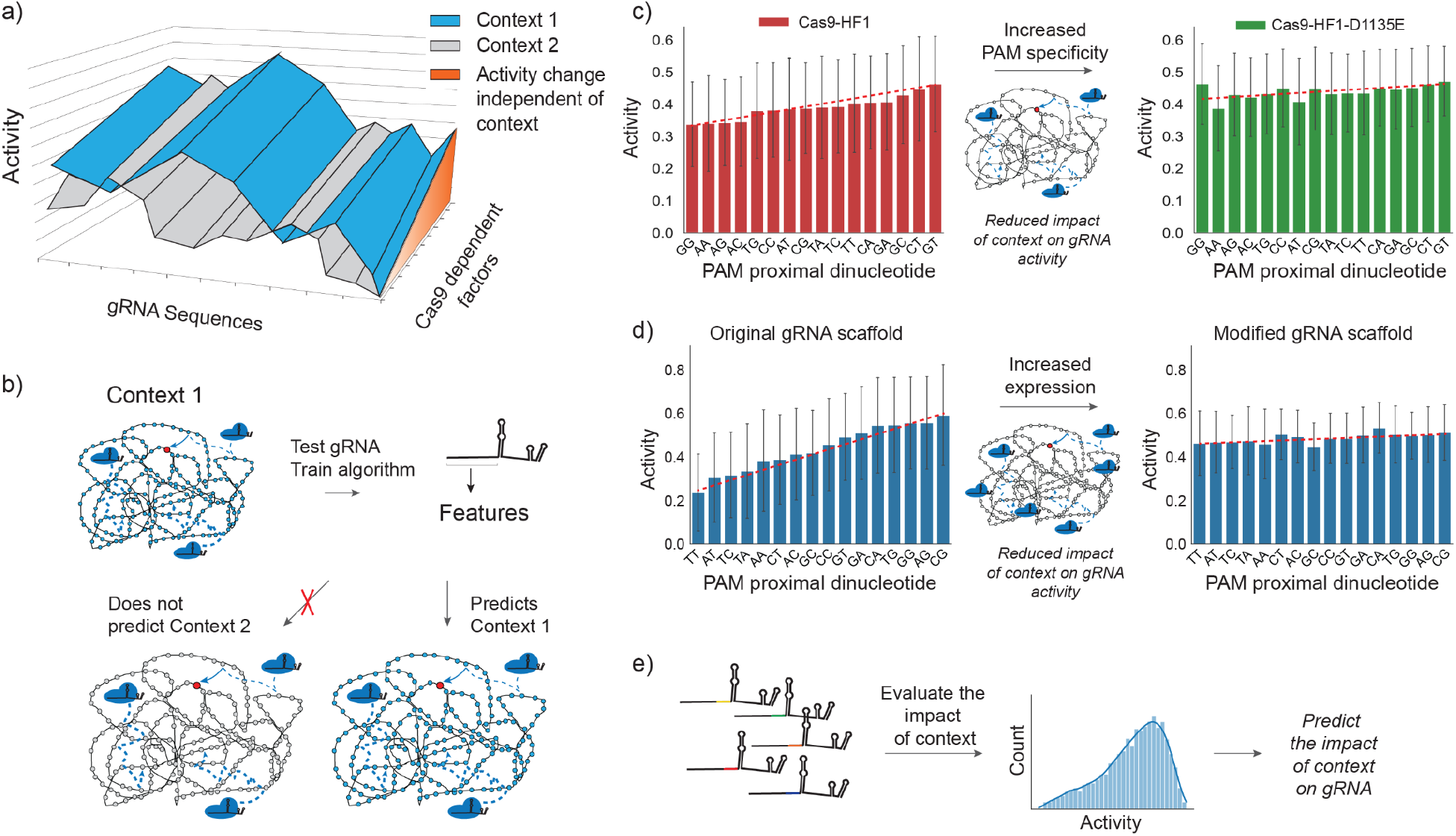
Cas9 activity is highly dependent on context. a) A given gRNA can be expected to have different activity in different host organisms in a sequence specific manner. Cas9 dependent factors that are independent of host context present an orthogonal axis of activity. b) Therefore, current algorithms trained on gRNA sequence features can perform well within the same context but will not accurately predict other species. However, the impact of context on gRNA activity can be reduced through c) increasing the specificity of Cas9 PAM binding to reduce potential interactions at non-target sites and d) increasing the expression of Cas9 and/or gRNA. e) Understanding the PAM proximal impact of context on gRNA activity also allows targeted gRNA libraries to specifically evaluate context effects (see Supplementary discussion on evaluating context).

This analysis also highlights factors that mitigate the impact of context on activity and helps to explain differences observed in many studies. In particular, high expression levels of Cas9 and/or gRNAs can reduce the impact of the genomic context on gRNA activity, improving on-target activity (Figure 6d). However, this may not be a general solution as high expression levels are also correlated with increased off-target activity.^30,31^ In cases where it is important to avoid off-target activity, other strategies may be preferred. One such strategy is to use Cas9 variants with higher PAM specificity (such as the D1135E mutant^27^), thus limiting the inhibitory non-target pool (Figure 6c).^16,27,32^ Higher PAM specificity mutants may be particularly useful in host contexts where host specific predictive algorithms have not yet been developed.

In addition to strategies for improving Cas9 activity in different contexts, this analysis emphasizes that many factors may negatively influence Cas9 activity relative to a maximal activity predicted by the PAM proximal sequence. While some of these factors, such as unwanted secondary structure in the gRNA or Cas9 preference for NGGH, are known, there is still much to learn.^3,33^ For example, several reports have highlighted specific motifs or nucleotide preferences of high fidelity variants but mechanistic explanations for this are lacking.^8,20,24,26^ Additionally, other contextual factors such as target site accessibility or other unknown chromosomal factors may play a role in Cas9 activity.^7,8,34^

Understanding how context impacts on-target activity may also help elucidate factors impacting off-target activity. Our analysis suggests that on-target activity is driven by factors outside of the target site. Since these contextual factors impact activity in a gRNA sequence-dependent manner, they are likely also relevant to off-target activity in the same way. Similarly, other unknown contextual factors may contribute to apparent sequence-dependence of off-target mismatch tolerance.^30^

Finally, while much of the current understanding of Cas9 activity has been limited to a perspective focused on the target site, it may be equally important to understand sequence specific differences at interactions at non-target sites as it is our view that the sum of these transient interactions is likely one main driver of context dependent differences. Several reports have found moderate or no connection between the number of predicted off-target sites and on-target activity.^10,35^ However, off-target sites make up a small minority of the potential search space when including transient non-target interactions.^16,36^ To our knowledge, transient interactions have only been evaluated in a handful of studies and no direct comparison of sequence dependent effects has been reported to date.^16,36^ *In vitro* studies by Sternberg *et al* demonstrated that Cas9 spent 1/10th the amount of time interrogating non-target sites with a 4 bp match than it did with non-target sites containing an 8 bp match.^36^ However, for any given gRNA there are likely to be orders of magnitude more non-targets with 4 bp matches than with 8 bp matches, without accounting for possible mismatches. This suggests that understanding transient interactions may be crucial to developing a better understanding of the sequence features driving context dependent differences. In the future, an improved understanding of context may well lead to 1) improved algorithms for predicting gRNA activity in established and novel organisms, 2) Cas9 variants with improved on-target and reduced off-target cleavage, 3) improved high-throughput functional screens, and 4) a better understanding of the factors driving activity in next generation CRISPR applications.

## Acknowledgements

We would like to acknowledge the following support: ONR YIP #12043956, and DOE EERE grant #EE0007563. We would also like to acknowledge support from Duke Innovation & Entrepreneurship Initiative.

## Author contributions

E.A. Moreb performed computational analyses. E.A. Moreb and M.D. Lynch designed analyses, analyzed results, wrote, revised and edited the manuscript.

## Methods

### Compiling datasets

We compiled 39 datasets from 19 papers (an overview is provided in Supplementary File 1, while datasets grouped by species are provided in Supplementary Files 2-6).^3–11,16,18–26^ We first filtered the data, only including results for gRNA where 1) we could find a matching target site in the target genome (if targeting an endogenous site) and 2) gRNA targeting NGG PAM sites. The following reference genomes were used: hg38 for human datasets (GenBank: GCA_000001405.15),^37^ mm9 for mouse datasets (GenBank: GCA_000001635.1),^38^ danRer10 for zebrafish (GenBank: GCA_000002035.3),^39^ W29 for *Y. lipolytica* (GenBank: GCA_003054345.1),^40^ MG1655 (GenBank: U00096.2)^41^, BW25113 (GenBank: CP009273.1)^42^ and W (GenBank: GCA_000184185.1)^43^ for *E. coli*. We report activity as it was reported in the original dataset but have inverted the sign on several datasets to ensure that in our comparisons more positive numbers correlate with more active gRNA. Datasets for which we inverted the sign include Xu *et al* 2015^6^, Moreb *et al* 2017^22^, Schwartz *et al* 2019^11^, Moreb *et al* 2020^16^, and Park *et al* 2021^23^. The data in Supplementary Files 2-6 include this sign inversion. When plotting datasets together (as done in Figures 4 and 5), we have re-scaled the activity measurements to values between 0 and 1, where 1 represents the most active gRNA.

For several datasets, we only used a subset of the available data. From Hart *et al* 2016^4^, for example, we only used the data from the Hct116 cell line, as described by Haeussler *et al* 2016^15^. This dataset included 4239 gRNA with activity averaged over several time points from 8 to 18 days.^4,15^ From Wang *et al* 2014^2^, we took data for cell lines KBM7 and HL60 that targeted essential genes, as provided by Xu *et al* 2015^6^. For datasets from Kim *et al* 2020a^19^, we only included gRNA Library B from the data provided at lentiviral MOI of 5 and only included gRNA targeting lentiviral sites. Similarly for Kim *et al* 2020b^20^, we only took data from gRNA Library B and we excluded repeat gRNA. From Park *et al* 2021^23^, we only took data from Library 1. Schwartz *et al* 2019^11^ performed library experiments in the presence and absence of the native NHEJ repair pathway. We used the Cutting Score results in the absence of NHEJ as this was not dependent on gene disruption by indels and therefore provided a more accurate measure of Cas9 activity.^11^ In addition to the data collected in mouse and human cell lines in their lab, Doench *et al* 2014^3^ provide data extracted from Shalem *et al* 2014^44^ of gRNA targeting essential genes. Finally, for Tálas *et al* 2021^24^ we combined the “balanced” datasets provided by the authors as a subset of the larger ~1.2 million gRNA library. The “balanced” datasets were provided by the authors to better help differentiate features that drive differences in efficient and inefficient gRNA as the majority of the larger ~1.2 million gRNA library would be deemed efficient.

### Assessing the importance of previously reported sequence features

We collected features specifically mentioned in the main text of papers as we reasoned this represents the features the authors deemed most important for activity. For each feature we determined if it was a discrete feature (ie, guanine in position 20 of the gRNA) or continuous feature (ie, GC content). To determine if a discrete feature positively or negatively impacted gRNA activity in a specific dataset, we calculated a log odds ratio based on the frequency of said feature in the most active third of gRNA versus frequency in the least active third of gRNA. If the log odds ratio was negative, the feature was said to negatively impact gRNA activity and if it was positive it would be described as positively impacting activity. For continuous features, we used a Pearson correlation with gRNA activity to determine if the relative impact of a feature was positive or negative based on the sign of the correlation. A feature would be considered to be in agreement across all datasets if the sign of the log odds ratio or Pearson agreed across all datasets, indicating a uniformly positive or negative impact on gRNA activity. Data is compiled in Supplemental File 7.

### Computational analyses

All computation was performed in Python with standard libraries, including: Datasets were managed with Pandas,^45^ NumPy was used for calculations,^46^ Regex was used for finding gRNA sequences in reference genomes,^47^ Scipy was used for statistics,^48^ and scikit-learn was used for linear regressions.^49^ Seaborn and Matplotlib were used for plotting.^50,51^ Biopython was used for calculating melting temperatures.^52^ Folding energies of gRNA were calculated using ViennaRNA RNAfold package.^53^ All code is provided in a Jupyter Notebook in Supplementary File 8.

