## Supplementary Materials for "An Analysis of gRNA Sequence Dependent Cleavage Highlights the Importance of Genomic Context on CRISPR-Cas Activity"

Moreb, E.A[1], and Lynch, M.D.[1,2,3]

[1]Department of Biomedical Engineering, Duke University

[2] To whom all correspondence should be addressed.

[3]

### Supplemental Materials

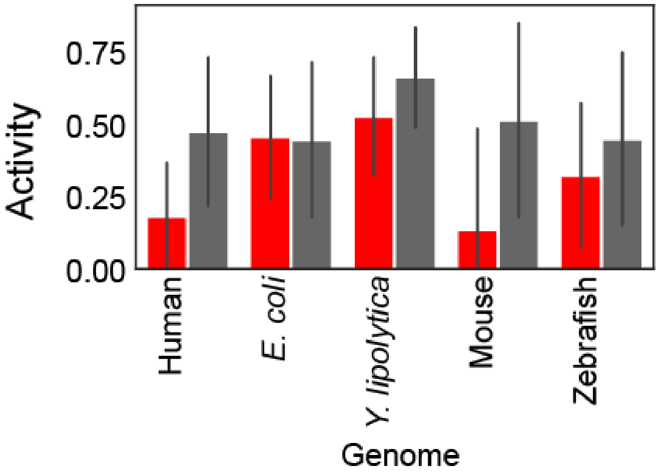


**Figure S1**: Average activity of gRNA containing four thymines (red) anywhere in the 20 bp targeting portion of the gRNA vs average activity of gRNA without four thymines (grey).

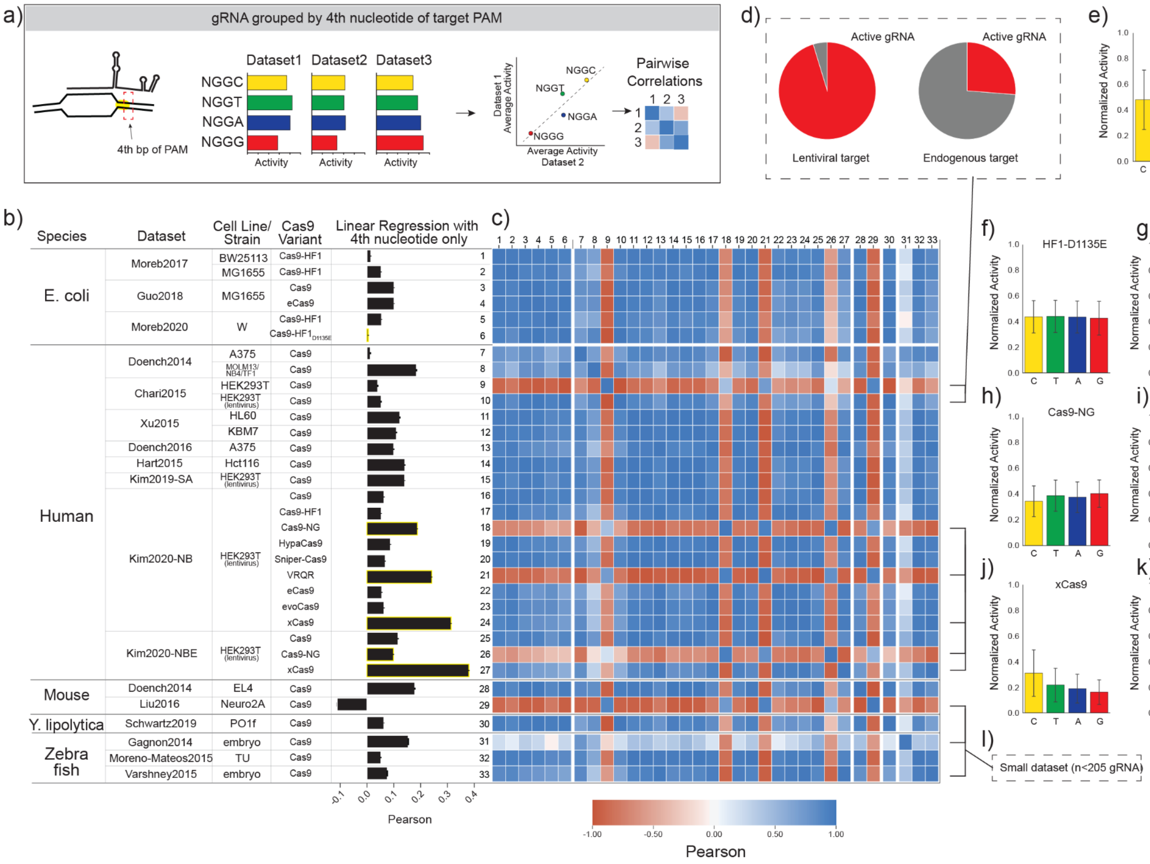

**Figure S2**: Validation of Cas9 PAM preference across datasets. a) Cas9 has been reported to have a slight preference for NGGH PAM sites, where H is either A, C, or T. We grouped gRNA by the 4th nucleotide of the PAM site, calculated the average activity per group, and correlated the averaged values between datasets in a pairwise fashion. b) We also correlated activity within each dataset with the 4th nucleotide of the PAM using a linear regression. Cas9 variants with mutations in the PAM interacting domain are highlighted in yellow. c) Pairwise correlations between all datasets show strong agreement in relative activity, confirming the preference for NGGH. Some datasets show weaker or no correlations. d) In Chari *et al* 2015, most of the gRNA targeting endogenous sites showed no activity, overriding any impact that PAM preference may have. e) Excluding the outlier datasets and Cas9 variants with mutations in the PAM interacting domain (f-k), the averaged values for all datasets confirms Cas9 preference for NGGH is consistent across different contexts. l) Smaller datasets with 205 gRNA or less were less consistent in capturing the Cas9 PAM preference. Among all datasets, three datasets did not have variability in the 4th nucleotide of the PAM and were therefore excluded from this analysis: Wang *et al* 2019, Tálas *et al* 2021, and Park *et al* 2021.

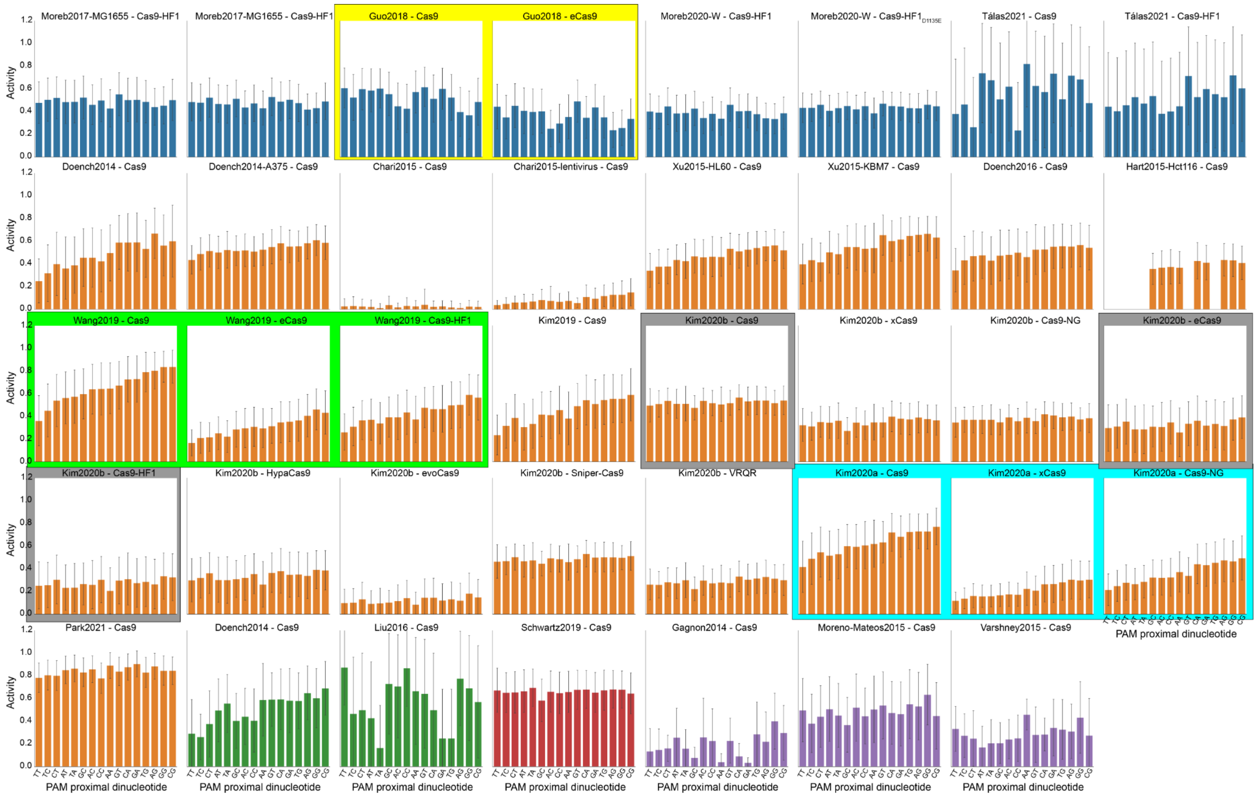


**Figure S3**: Activity of gRNA grouped by PAM proximal dinucleotide sequence. Plots represent the data in the pairwise correlations in Figure 4 in the main text. The dinucleotide sequences are ordered by least active to most active based on the largest human dataset, from Wang *et al* 2019[1]. Colors of bars indicate species groupings: Blue = *E. coli*; Orange = Human; Green = Mouse; Red = *Y. lipolytica*; Purple = Zebrafish. Additionally, for datasets where Cas9, Cas9-HF1, and/or eCas9 are reported together, we highlight that activity grouped by the PAM proximal dinucleotides does not significantly change. These data suggest that context specific impact on the PAM proximal sequence is consistent across these variants in data from Guo *et al* 2018[2] (yellow shading), Wang *et al* 2019[1] (green shading), Kim *et al* 2020b[3] (grey shading), and Kim *et al* 2020a[4] (teal shading).

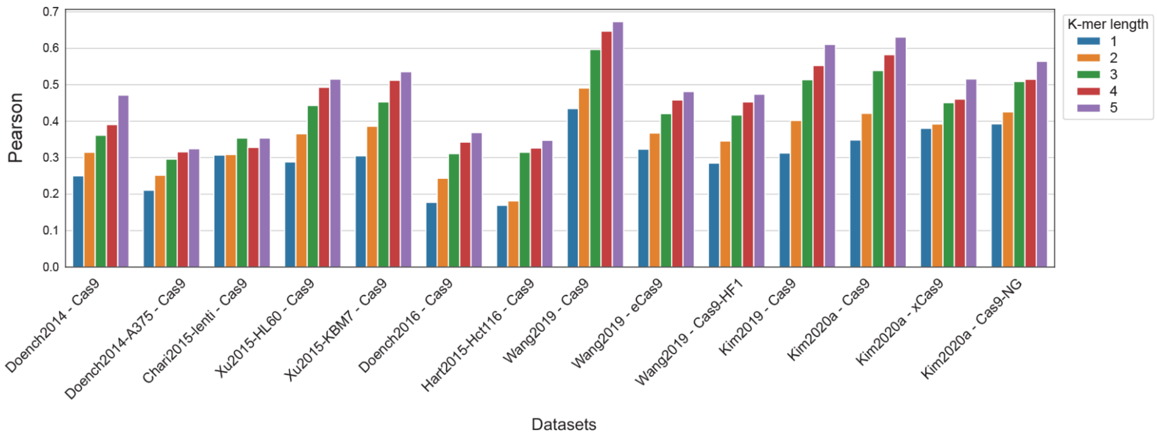

**Figure S4**: Predictions within human datasets using the PAM proximal sequence based activity averages (see Figure 5), improve with longer K-mer sequence. Excluded datasets with modified gRNA scaffold and the Chari *et al* 2015 dataset targeting endogenous sites, due to the overall low activity of that dataset.

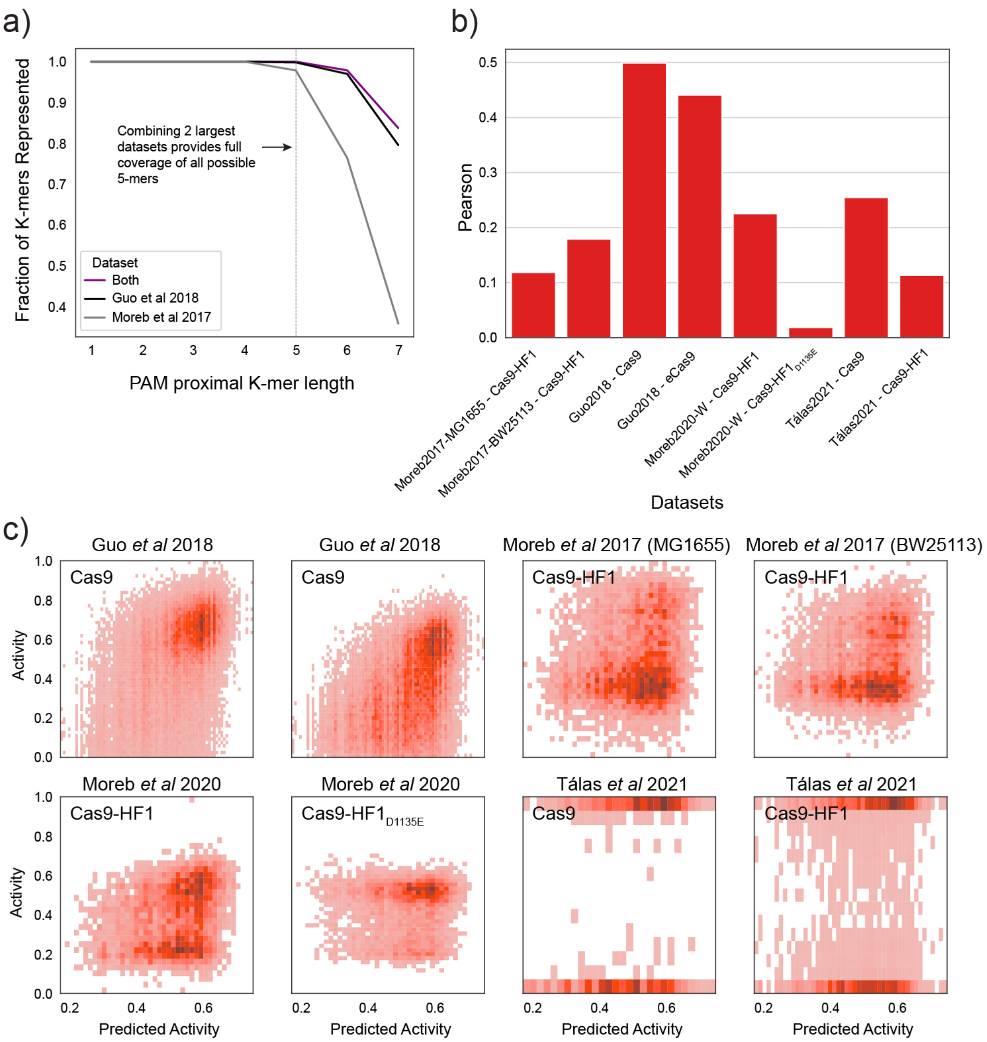


**Figure S5**: Using a similar approach as that of Figure 5 in the main text, we combined data from Guo *et al* 2018 and Moreb *et al* 2017 to produce predictions based on the PAM proximal 5 bp sequence. a) Combining these two datasets gives full coverage of all 5-mer sequences. b) Predictions based on the PAM proximal sequence are evaluated based on Pearson correlation with actual activity. c) For each dataset, we plot actual activity against predicted activity.

### Supplementary Note 1:

#### Proposed gRNA library guidelines for understanding context specific sequence preferences in new species

In this analysis, we highlighted that the PAM proximal sequence is most correlated with gRNA activity in a host specific manner, therefore it is important for predicting the activity of given gRNA within a specific context. In Figure S6, we show that increasing the length of the PAM proximal sequence used for predictions improves the predictive power of this approach. However, no datasets have yet been designed to systematically evaluate context dependent sequence-preference in this manner. For understanding context-dependent gRNA activity in new hosts, it may therefore be beneficial to systematically assess context dependent factors. Here we highlight factors to consider while designing a gRNA library in novel host contexts:

1. To systematically assess the impact of context, it may be beneficial to design sequence diversity in the PAM proximal sequence by selecting *n* replicates of each K-mer, where K represents the PAM proximal sequence length. In general, larger *n* and longer K will provide a better understanding of context. However, selecting the appropriate *n* and K values is dependent on practical variables, such as the throughput of the gRNA screen, budget, and constraints. To help researchers select the appropriate *n* and K for their gRNA library design, we calculated the library size with different *n* and K values (Figure S6a), as well as the potential predictive ability of a library designed with different *n* and K values (Figure S6b).
2. In addition to designing variability within the PAM proximal sequence, researchers may also want to account for unwanted secondary structure within the gRNA,[5,6] Cas9 preference for NGGH PAM sites,[7] and species-specific transcriptional modifiers (ie, four thymines in human or mouse cell lines[8]).

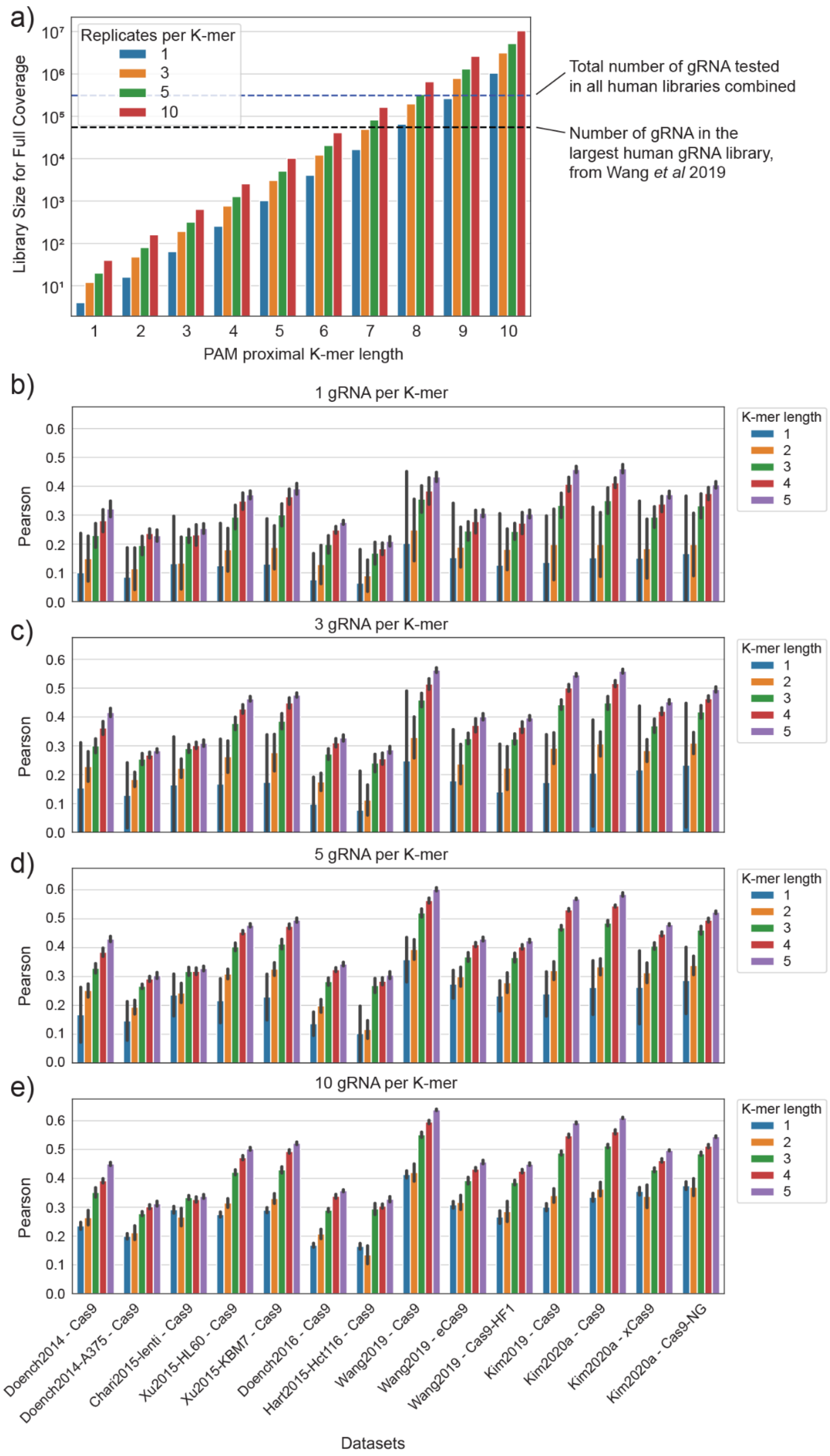


**Figure S6**: a) We calculate the required library size to provide complete coverage for all PAM proximal K-mer sequences (K between 1 and 10 base pairs) and for different numbers of replicates per unique K-mer. For example, to get complete coverage of all possible 5-mer PAM proximal sequences with 3 replicates per 5-mer, you would need a library size of just 3,072 gRNA. Provided as reference are the total number of gRNA in all the human datasets collected in this paper (black dashed line) and the number of gRNA in the largest dataset in this paper (blue dashed line). Following the steps of the analysis shown in Figure 5, we combined the wild-type Cas9 datasets from Wang *et al* 2019 and Kim *et al* 2019 and randomly sampled either b) 1, c) 3, d) 5, or e) 10 gRNA per K-mer to look at the number of replicates required to evaluate context specific differences in host context. After random sampling, we grouped gRNA by the PAM proximal K-mer sequence and used the averages to predict activity in other human datasets using the original gRNA scaffold. We repeated the sampling and prediction steps 10 times and plot the average of these 10 predictions.
